# Chemogenetic inhibition of a monosynaptic projection from the basolateral amygdala to the ventral hippocampus selectively reduces appetitive, but not consummatory, alcohol drinking-related behaviors

**DOI:** 10.1101/529719

**Authors:** Eva C. Bach, Sarah E. Ewin, Chelcie F. Heaney, Hannah N. Carlson, Antoine G. Almonte, Ann M. Chappell, Kimberly F. Raab-Graham, Jeffrey L. Weiner

**Affiliations:** Wake Forest School of Medicine, Department of Physiology and Pharmacology, Winston-Salem, NC 27157

**Keywords:** alcohol use disorder, anxiety, motivation, seeking

## Abstract

Alcohol use disorder (AUD) and anxiety/stressor disorders frequently co-occur and this dual diagnosis represents a major health and economic problem worldwide. The basolateral amygdala (BLA) is a key brain region that is known to contribute to the etiology of both disorders. Although many studies have implicated BLA hyperexcitability in the pathogenesis of AUD and comorbid conditions, relatively little is known about the specific efferent projections from this brain region that contribute to these disorders. Recent optogenetic studies have shown that the BLA sends a strong monosynaptic excitatory projection to the ventral hippocampus (vHC) and that this circuit modulates anxiety- and fear-related behaviors. However, it is not known if this pathway influences alcohol drinking-related behaviors. Here, we employed a rodent operant drinking regimen that procedurally separates appetitive (e.g. seeking) and consummatory (e.g. intake) behaviors, chemogenetics, and brain region-specific microinjections, to determine if BLA-vHC circuitry influences alcohol drinking-related measures. We first confirmed prior optogenetic findings that silencing this circuit reduced anxiety-like behaviors on the elevated plus-maze. We then demonstrated that inhibiting the BLA-vHC pathway significantly reduced appetitive alcohol drinking-related behaviors while having no effect on consummatory measures. Sucrose seeking measures were also reduced following chemogenetic inhibition of this circuit. Taken together, these findings provide the first indication that the BLA-vHC circuit may regulate appetitive alcohol drinking-related behaviors and add to a growing body of evidence suggesting that dysregulation of this pathway may contribute to the pathophysiology of AUD and anxiety/stressor-related disorders.

**HIGHLIGHTS:**

- The basolateral amygdala sends a monosynaptic glutamatergic projection to the ventral hippocampus
- Inhibiting this circuit reduces anxiety-like behaviors in male Long Evans rats
- Inhibition of this pathway also decreases operant alcohol seeking-related behaviors

## 1.0 INTRODUCTION

Alcohol use and misuse represent major health and socioeconomic concerns in the United States and around the world. A recent report found that nearly 7% of all global deaths could be attributed to alcohol and the 2016 Surgeon General’s Report on Alcohol, Drugs and Health noted that alcohol contributes to 88,000 (Sohi *et al*., 2021) deaths each year in the United States at a cost of almost $250 billion to our economy (Stahre *et al*., 2014; Sacks *et al*., 2015; U.S. Department of Health and Human Services (HHS), 2016). Despite these devastating numbers, some populations are particularly vulnerable to alcohol’s deleterious health effects. One notable group includes individuals diagnosed with anxiety and stressor-related disorders, like generalized anxiety disorder and post-traumatic stress disorder (PTSD). For example, 40-60% of individuals with PTSD also suffer from alcohol use disorder (AUD) (Shorter *et al*., 2015; Gilpin & Weiner, 2017; Petrakis & Simpson, 2017). Moreover, these comorbidities are associated with greater symptom severity for each disorder and a much poorer prognosis in recovery (Schafer & Najavits, 2007; Evren *et al*., 2011; Ipser *et al*., 2015). Importantly, despite the high prevalence of these comorbid conditions, there is marked variability in the likelihood of developing these disorders (Marmar *et al*., 2015; Walker *et al*., 2017) and very little is known about the specific neural circuits that might confer vulnerability or resilience to these diseases (Gilpin & Weiner, 2017; Carlson & Weiner, 2021).

One possible reason for the frequent co-occurrence of these disorders is that their etiology may involve maladaptive changes in common neural pathways. For example, many studies have reported that both AUD and PTSD are associated with dysregulated activity and connectivity in circuits that include the prefrontal cortex, amygdala, and hippocampus (Almonte *et al*., 2017; Skelly *et al*., 2017; Bloodgood *et al*., 2018; Santos *et al*., 2019; Carlson & Weiner, 2021). Notably, recent studies have demonstrated that these brain regions are not homogenous structures but, rather, are comprised of distinct populations of cells with unique afferent and efferent connectivity that often encode behaviors with opposing valence (Kim *et al*., 2016; Beyeler *et al*., 2018). A better understanding of the pathophysiology of AUD and comorbid disorders will require clear insight into the specific circuits that drive the maladaptive behaviors associated with these diseases.

One circuit that has received considerable attention of late involves a monosynaptic projection from glutamatergic pyramidal cells in the basolateral amygdala (BLA) to the ventral hippocampus (vHC). Recent optogenetic studies have demonstrated that activation of this circuit increases unconditioned anxiety-like behaviors and reduces social interaction in mice and that silencing this pathway promotes anxiolysis and disrupts foot shock, but not contextual, fear learning in rats (Huff *et al*.,2016). Moreover, we have recently demonstrated that withdrawal from chronic ethanol vapor exposure increases BLA-vHC synaptic excitability (Bach *et al*.,2021a). Despite the importance of this pathway in a number of behaviors associated with AUD and comorbid conditions, whether this circuit modulates alcohol drinking-related measures is not known. To that end, we integrated Designer Receptors Exclusively Activated by Designer Drugs (DREADDs) and brain region-specific microinjection methods to assess the effect of chemogenetic inhibition of the BLA-vHC circuit on alcohol drinking-related behaviors using a limited-access operant regimen that procedurally separates appetitive (seeking) and consummatory drinking measures.

## 2. METHODS

### 2.1 Subjects

Four separate cohorts of 8 male, Long Evans rats were used in these studies, all obtained from Envigo (Indianapolis, IN). Animals arrived at 150-175g, and were singly housed in clear Plexiglas cages (25.4 cm × 45.7 cm). Rats were weighed and handled daily. Rats had *ad libitum* access to food (Prolab RMH 3000, LabDiet; PMI Nutrition International, St. Louis, MO) and water throughout and were maintained on a 12-h light/dark cycle in the same colony room. All behavioral experiments were conducted between 6-10 a.m. (EST), 5 days/week. Animal care procedures were carried out in accordance with the NIH Guide for the Care and Use of Laboratory Animals and were approved by the Wake Forest University Animal Care and Use Committee.

### 2.2 Surgery and Microinjection Procedures

Animals were anesthetized with sodium pentobarbital (50mg/kg intraperitoneally) and placed into a rodent stereotaxic instrument. After exposure of the skull, a viral construct expression the inhibitory DREADD, hM4D(Gi) (AAV5-CamKIIa-hM4D(Gi)-mCherry (Gi-DREADD; UNC Vector Core, Chapel Hill, NC or Addgene) or the reporter protein EGFP (AAV5-CaMKIIa-EGFP (UNC Vector Core, Chapel Hill, NC) was bilaterally injected in the BLA (coordinates: anteroposterior −2.8 from bregma; mediolateral +5.0 mm, and ventral from the top of the brain −7.3 mm) for all behavioral/histological studies. Following this, stainless-steel guide cannulas (26 gauge) were implanted to terminate 1mm above the vHC (coordinates: anteroposterior −5.6 mm from bregma; mediolateral from the midline +5.0 mm; ventral from the top of the brain –5.0mm). Using stainless-steel screws and dental cement, the guide cannulas were attached to the skull. After 3-4 weeks of viral incubation, microinjections began. Animals were placed in a small plastic container and restrained by hand while the obturators were removed and injectors were inserted. The injectors extended 1 mm beyond the end of the guide cannula and the drug solution (aCSF or CNO dissolved in aCSF at 10ng, 33ng or 100ng per side in a Latin Square design) was bilaterally infused for 1 min (0.5μL per side). The dose range selected for CNO injections was based on a prior report that local microinjection of 50 ng CNO into the dorsal striatum could influence goal-directed behavior (Gremel et al., 2016). Following the one minuteinjections, the injectors were left in place for an additional 30 seconds. Five minutes following the microinjections, animals were placed in the elevated plus-maze or operant chambers tobegin behavioral testing. For electrophysiological confirmation of CNO-dependent DREADD terminal inhibition the a viral construct expressing the inhibitory DREADD, hM4D(Gi) (AAV5-CamKIIa-hM4D(Gi)-mCherry (Gi-DREADD; UNC Vector Core, Chapel Hill, NC) and the excitatory opsin, Channelrhodopsin (AAV5-CaMKIIa-hChR2(H134R)-EYFP (Addgene, Cambridge, MA) was bilaterally injected in the BLA or aBLA (coordinates: anteroposterior −1.96 from bregma; mediolateral +5.0 mm, and ventral from the top of the brain −8.15 mm).

### 2.3 Histology and imaging

One cohort of animals (n=8; chemogenetic ethanol operant self-administration cohort) was processed for detailed histological determination of viral spread throughout the BLA and the vHC. Animals were deeply anesthetized using Euthanasia solution (source) and transcardially perfused with phosphate buffered saline (PBS) followed by 4% paraformaldehyde (PFA) for approximately 20 minutes per solution. The brains were then removed and immediately placed in 4% PFA where they were stored overnight. Brains were subsequently washed with PBS and placed into a solution containing 30% sucrose in PBS until they fully sunk to the bottom of their container (~1 week). Transverse slices (25 μm) containing the BLA or vHC were then cut on a Leica SM2010F sliding microtome and placed in cryoprotectant (10% glycerol, 15% ethylene glycol in 0.05 M PBS) until further processing. For staining, slices were washed with PBS three times for 5 min each, stained in DAPI (30 nM; Invitrogen) for 30 min and subsequently washed again in PBS three times for 5 min. Stained slices (~200 μm apart) throughout the BLA and vHC were mounted and cover slipped. All images were acquired on a Nikon A1plus confocal microscope using the same acquisition settings. Slices were imaged using an air 20× lens at 512×512 pixels in a given Z plane where the focus was based on virus staining. Large images were obtained using 10% overlap stitching. Images were post-processed to enhance visualization. To construct maps of spread around the BLA injection site (8 rats) and the spread of projections in the vHC (6 rats) the spread from each animal and hemisphere (bilateral spread) was transposed onto a single hemisphere at multiple stereotaxic sites (BLA-9 sites, vHC-8 sites) from bregma. In regions where cannula damage interfered with a determining viral expression at the site of the cannula, viral expression through this region was extrapolated from the expression seen in undamaged regions immediately caudal and rostral to it. In 3 animals viral expression could only be detected within a single hemisphere. Cannula placements were visually confirmed following transcardial perfusions and sectioning in both operant self-administration (ethanol and sucrose) cohorts for which data is presented (n=15).

### 2.4 Slice Electrophysiology

Animals were deeply anesthetized using isoflurane. Following decapitation, the brain was removed rapidly and suspended in ice-cold NMDG recovery solution containing (in mM): 92 NMDG, 2.5 KCl, 1.25 NaH_2_PO_4_, 30 NaHCO_3_, 20 HEPES, 25 glucose, 2 thiourea, 5 Na-ascorbate, 3 Na-pyruvate, 0.5 CaCl_2_·H_2_O, and 10 MgSO_4_·H_2_O. NMDG was titrated to pH 7.4 with 17 mL +/− 0.5 mL of 5 M hydrochloric acid. Transverse slices containing the vHC were cut at a thickness of 300 μm using a VT1000S Vibratome (Leica Microsystems). vHC slices were placed in a holding chamber containing NMDG recovery solution. Slices were allowed to recover for 35 min before being transferred to a chamber containing artificial cerebral spinal fluid (aCSF) containing (in mM): 125 NaCl, 1.25 NaH_2_PO_4_, 25 NaHCO_3_, 10 D-Glucose, 2.5 KCl, 1 MgCl_2_, and 2 CaCl_2_. NMDG recovery and aCSF holding solutions were oxygenated with 95% O_2_ and 5% and warmed to 32-34 °C. For recordings, a single brain slice was transferred to a chamber mounted on a fixed stage under an upright microscope (Scientifica SliceScope Pro 2000 microscope), where it was superfused continuously with warmed oxygenated aCSF. Whole-cell voltage-clamp recordings were made from presumptive pyramidal neurons of the ventral hippocampus (vHC) using recording pipettes pulled from borosilicate glass (open tip resistance of 6–9 MΩ; King Precision Glass Co., Claremont, CA). The pipette solution contained (in mM): 130–140 Cs-gluconate, 10 HEPES, 1 NaCl, 1 CaCl_2_, 3 CsOH, 5 EGTA, 2 Mg^2+^-ATP, 0.3 GTP-Na_2_ and 2 Qx-314. Intracellular Cs^+^ was used as the primary cation carrier in voltage-clamp recordings to block K^+^ currents, including postsynaptic GABA_B_ receptor-mediated currents, in the recorded neuron. Neurons were targeted for recording under a 40x water-immersion objective (numerical aperture = 0.8) with infrared-differential interference contrast (IR-DIC) optics, as described previously. Electrophysiological signals were obtained using a Multiclamp 700B amplifier (Molecular Devices, Union City, CA), low-pass filtered at 2 or 3 kHz, digitized at 10kHz, and recorded onto a computer (Digidata 1440A, Molecular Devices) using pClamp 11.0 software (Molecular Devices). Seal resistance was typically 2–5 GΩ and series resistance, measured from brief voltage steps applied through the recording pipette (5 mV, 5 ms), was <25 MΩ and was monitored periodically during the recording. Recordings were discarded if series resistance changed by >20% over the course of the experiment. Recordings were made from six rats and each recorded neuron represented an individual data point (n). Optical stimulation was performed using blue light (473 nm) to stimulate terminals originating from the BLA synapsing onto vHC pyramidal neurons. Recordings were conducted in the presence of picrotoxin (100 μM) at a holding potential of −70 mV. The amplitudes of a minimum of 10 optically evoked excitatory postsynaptic currents (oEPSCs) were averaged to establish the synaptic input under baseline conditions. CNO was bath applied for a minimum of 10 min to establish the CNO-dependent inhibitory effect on glutamatergic BLA-driven inputs.

### 2.5 Behavioral Tasks

#### 2.5.1 Operant Self-Administration

Daily sessions were performed in commercially available, sound-attenuated operant chambers (Med Associates, East Fairfield, VT) as previously described (Samson et al., 1999). Each chamber contained a house light to signal the beginning of a session, a retractable leverand a sipper tube that extended into the chamber. The system was computer controlled and appetitive and consummatory responses were collected at 2 Hz and analyzed using MedPC software (Med Associates, St. Albans, VT).

To train rats to self-administer alcohol (ethanol), we used an abbreviated sucrose-substitution protocol based on the method of Samson (Samson, 1986). Briefly, on the first day of shaping, animals were acclimated to chambers for two hours using a 10% sucrose solution on a fixed ratio (FR1) schedule that resulted in 45 seconds of access to the sipper. Over the next consecutive days, session time, fixed ratio, and sipper access time were modified, and the sucrose concentration was decreased by 1% each day and replaced with increasing concentrations of ethanol. The fixed ratio schedule was increased from an FR1 to an FR4 over an 8-day period. On day 9, the schedule was changed such that completion of a response requirement of 8 lever presses (RR8) resulted in a twenty-minute presentation of the reinforcer. Over the next 10 sessions, the response requirement was increased to an RR30 and the final solution used in testing was 10% ethanol. A similar protocol was also used to train rats to an RR30 for a 3% sucrose solution. Animals had 20 minutes to complete the RR30, although subjects typically completed the lever press response requirement in less than five minutes. Appetitive and consummatory behaviors were monitored daily and data was collected using Med Associates software. Consummatory measures included ethanol intake, latency to first lick, lick rate, and lick bouts, with a bout being defined as a licking episode without any pause greater than 20 seconds. Appetitive measures included latency to first lever press, lever press rate, and lever press bouts, defined as a lever press episode without any pause greater than 20 seconds.

#### 2.5.2 Microinjections

After 3-4 weeks of viral incubation, and continual sessions in the operant chambers, (5 days/week, Mon-Fri), microinjections began using the timeline illustrated in Fig 1. Animals were placed in a small plastic container and restrained by hand while the obturators were removed and injectors were inserted prior to the start of the operant session microinjections. Microinjections were given once/week. All microinjections (CNO or saline) were injected at a rate of 0.5 μL/minute over the course of 1 minute. Animals were observed for an additional 6 minutes and subsequently placed into the operant chambers and allowed to respond to the lever for 20 minutes. Three doses of CNO (10 ng/side; 33 ng side, 100 ng/side) were tested on standard drinking sessions using a within-subjects, Latin Square design. Importantly, on microinjection days, all rats were given 20 minute access to the reinforcer solution following the 20 minute lever press time, regardless of whether they completed the RR30, in order to assess the effects of CNO injections on consummatory behaviors. Cohorts injected with the Gi-DREADD construct received 5 microinfusions of ACSF or CNO and the cohort injected with a construct that only expressed a reporter protein received 6 microinjections (this cohort was also tested on the elevated plus-maze).

**Figure 1.**
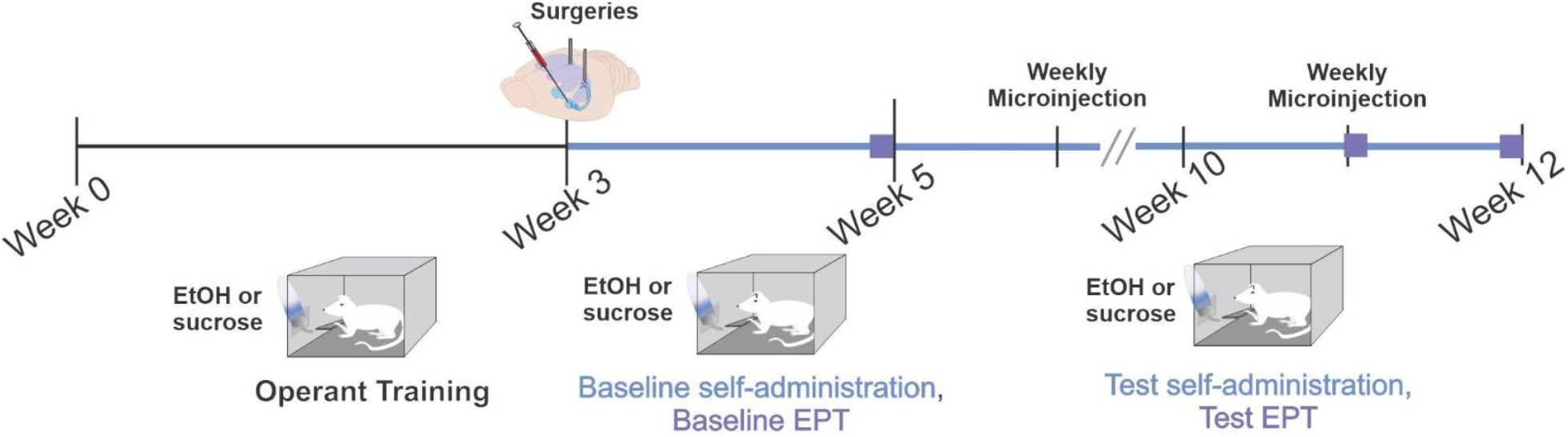
Schematic illustration of the timeline for chemogenetic microinjection studies.

#### 2.5.3 Extinction Probe Trials

To obtain an appetitive measure devoid of any consummatory behaviors, extinction probe trials were conducted. On the days when extinction probe trials were performed, rats were placed into the operant chamber and allowed to respond on the lever for the entire 20 minutes, regardless of the number of lever presses completed. Following this period, all subjects were presented with the reinforcer and allowed to drink for 20 minutes. Three extinction probe trials were conducted on each cohort, at least one week apart. The first was a baseline trial (no microinjections) and the other two trials followed intra-vHC microinfusion of CNO (100 ng/per side) or vehicle with order of testing controlled using a Latin-Square design.

#### 2.5.4 Elevated Plus-Maze

To measure unconditioned anxiety-like behavior, identical microinjection procedures were used to test behavioral responding on standard elevated plus-mazes (Med Associates, St.Albans, VT). The elevated plus-mazes were raised 72.4 cm from floor level, with runways measuring 10.2 cm wide by 50.8 cm long. Open runways had 1.3 cm high lips and closed runways were detected via infrared sensors attached to the opening of each arm of the maze. Data were obtained and recorded via a computer interfaced with control units and MED-PC programming (Med Associates). Animals received vehicle or CNO (100 ng per side) and, after five minutes, were placed at the junction of the four arms, facing one of the open arms and allowed to freely explore the maze for five minutes. Open arm time and open arm entries were used as measures of anxiety-like behavior and the number of closed arm entries was measured to assess general locomotor activity.

#### 2.5.5 CNO Control Cohorts

To control for non-specific effects of CNO, a separate cohort of 8 rats received intra-vHC microinjections and was tested on the elevated plus-maze and operant alcohol self-administration regimen described above. However, this cohort received BLA microinjections of a virus that only expressed a reporter protein (AAV5-CaMKIIa-EGFP). Seven of these subjects completed the complete alcohol self-administration procedure but only five completed both ACSF and CNO treatments on the elevated plus-maze (due to loss of head caps).

### 2.6 Blood Ethanol Determination

Blood ethanol concentrations were measured two days after the final self-administration microinjection session immediately following operant self-administration. A 10 μL blood sample was collected from a tail snip of each rat. Blood ethanol concentrations were determined using a commercially available ethanol dehydrogenase enzymatic assay kit (Diagnostic Chemicals, Oxford, CT).

### 2.7 Statistical Analysis

Statistical analyses were performed in SigmaPlot version 14.0 and GraphPad version 9.0. Normality was assessed using a Shapiro-Wilk Normality Test. Data were analyzed using two-tailed t tests or the Wilcoxon signed-rank test, one-way repeated measures ANOVAs, and mix model two-way ANOVAs. A p value of 0.05 or less was considered significant.

## 3.0 RESULTS

### 3.1 Immunohistochemical and electrophysiological confirmation of Gi-DREADD expression

Detailed visual confirmation of reporter protein expression and diffusion within both the BLA and vHC terminal fields was performed on a subset of 8 animals using immunohistochemistry for DAPI (nuclear marker, blue) and the viral construct reporter protein mCherry (red) visualized with confocal microscopy. To get a better assessment on the level of viral spread of transfection in cell bodies of the BLA and its terminal projections in the vHC, 25 μm slices ~200 μm apart from one another were imaged throughout the BLA and vHC. Viral transfection at the injection site was seen most robustly in the BLA, spreading along the anterior-posterior axis. Viral spread typically extended into to lateral amygdala. Outside of the amygdala, we commonly observed expression extending into the caudate putamen and the dorsal endopiriform nucleus. We also observed minor and less consistent expression in the piriform cortex and the central amygdala (Fig 2 A-C).

**Figure 2.**
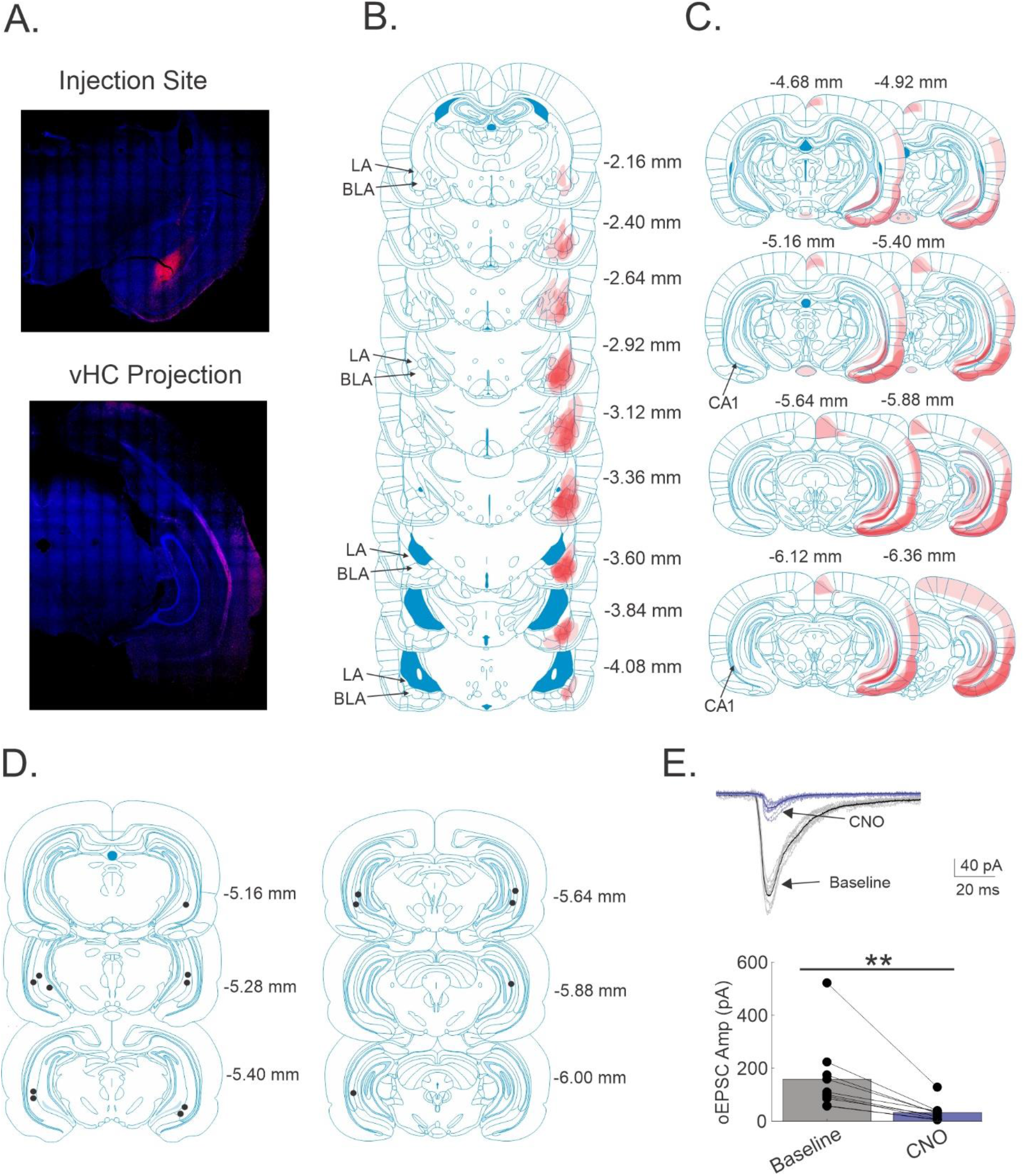
The BLA makes monosynaptic excitatory projections to the vHC that can be inhibited by CNO in animals expressing ChR2 and GiDREADD. A) Representative images of viral expression at the injection site (top panel) in the BLA and its projections in the vHC (lower panel). Nuclear DAPI staining is shown in Blue and viral expression is shown in red. B) Schematic illustration of viral spread and expression in and around the injection site. C) Schematic illustration of viral spread throughout the vHC. In (B) and (C) numbers to the right of each brain slice indicate anterior/posterior stereotaxic coordinates relative to bregma. To construct composite maps expression/projection spread in the target regions (BLA and vHC) from each animal and hemisphere was overlaid on a single hemisphere. Color gradient (from light to dark red) within injection/projection sites illustrates the consistency of viral expression/projection within the overall area among individual animals and hemispheres. D) Schematic illustration of cannula placements in the BLA. E) CNO inhibits optically evoked BLA inputs in the vHC of animals coexpressing ChR2 and GiDREADD (Baseline: 158.3 ± 48.8 pA, CNO: 32.5 ± 12.7 pA, n=9, Wilcoxon signed-rank test, **p<0.01). E) Representative oEPSCs traces (upper panel) in a vHC neuron during baseline (black traces) and CNO application (purple traces). Thin lines represent individual responses while thick lines illustrate average response during baseline and CNO application. Bar graph (lower panel) of oEPSC amplitudes at baseline and during the application of CNO illustrating responses for individual neurons (connected lines).

Viral spread in the vHC was seen in area CA1, the subiculum, as well as the entorhinal cortex, in line with previous reports identifying projections to these vHC subregions (Pikkarainen *et al*., 1999). In some instances we also identified viral spread into cortical regions of the sensory systems. All cannula placements were identified in the hippocampal regions (in or near CA1; Fig 2D) and any of the “off target” areas should not have been exposed to meaningful CNO concentrations to be pharmacologically impacted by the microinjections.

To confirm the CNO-dependent inhibitory effect at BLA terminals synapsing onto neurons in the vHC, we conducted whole-cell electrophysiology in acute brain slices from animals that were transfected with ChR2 and Gi-DREADD in the BLA. Neurons in the vHC were targeted for voltage-clamp recordings at −70 mV in the presence of the GABA_A_ receptor antagonist, picrotoxin, to isolate glutamate-dependent EPSCs. BLA terminals were optically stimulated and a baseline measure of oEPSC amplitudes was determined. Subsequent bath application of CNO resulted in significant decrease in oEPSC amplitudes (Baseline: 158.3 ± 48.8 pA, CNO: 32.5 ± 12.7 pA, n=9, Wilcoxon signed-rank test, p<0.01; Fig 2E). These results confirm that BLA terminal activation of the inhibitory Gi-DREADD is effective at inhibiting neurons of the vHC.

### 3.1 Chemogenetic inhibition of the BLA-vHC circuit reduces anxiety-like behaviors in the elevated plus-maze

To confirm prior optogenetic studies, we first tested the effect of silencing the BLA-vHC pathway on anxiety-like behaviors using the elevated plus-maze. In rats expressing hM4D(Gi)-DREADD, intra-vHC infusion of CNO (100 ng per side) significantly increased open arm time (t_1,12_=−3.445, p < 0.02; Fig 3A) and open arm entries (t_1,12_=−3.197, p<0.01; Fig 3A), two well-validated measures of anxiety-like behavior. Closed arm entries, a measure of non-specific locomotor activity in this assay, was not affected (t_1,12_=1.336, p>0.05; Fig 1A). To ensure that these anxiolytic effects were dependent on Gi-DREADD expression, we generated a separate cohort of rats that received intra-BLA infusion of a virus that only expressed a reporter protein. Intra-vHC infusion of CNO in these animals had no effect on open arm time (t_1,8_=1.131, p>0.05), open arm entries (t_1,8_ = 1.000, p>0.05) or closed arm entries (t_1,8_ = −0.894, p > 0.05) (Fig 3B).

**Figure 3.**
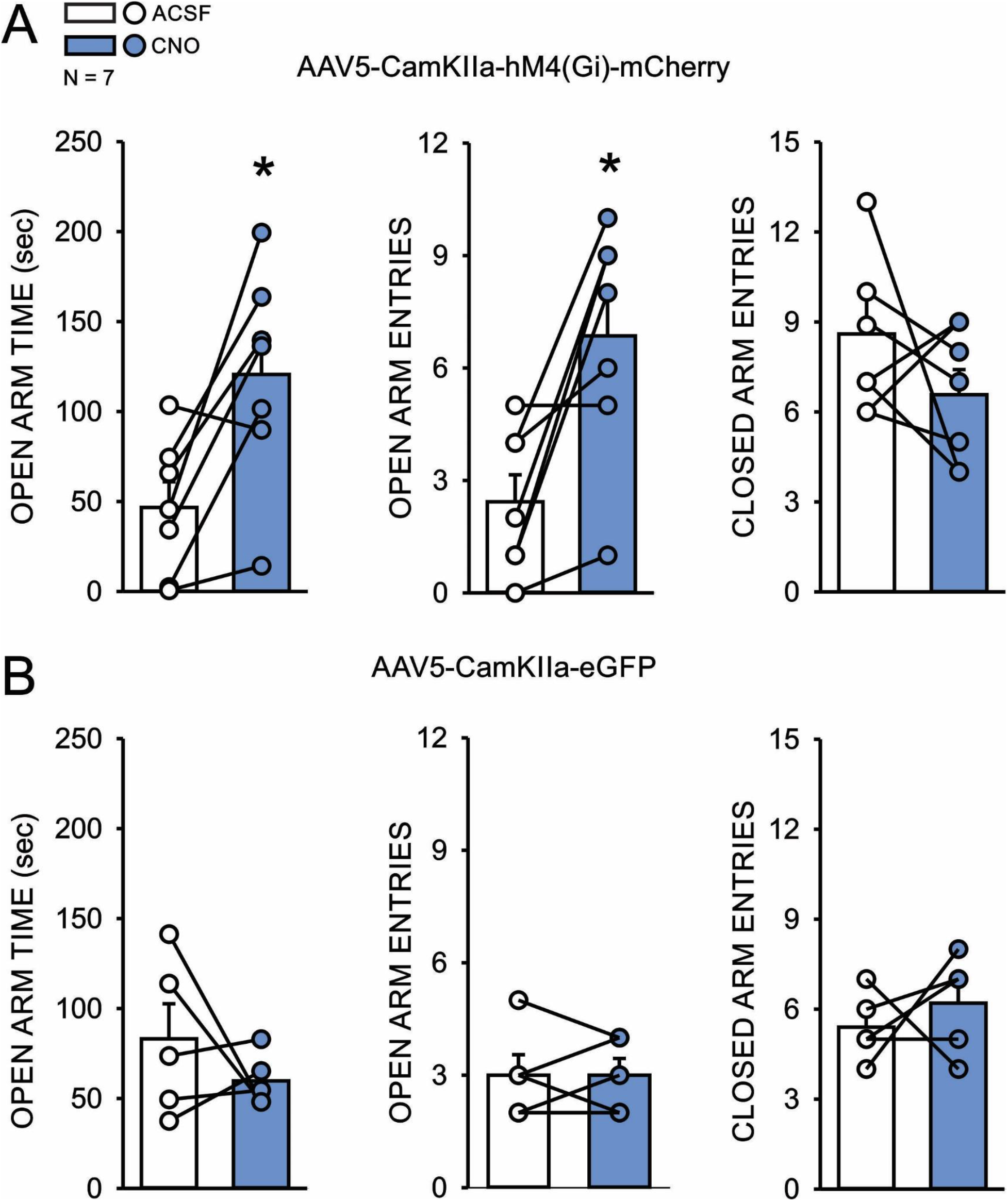
Chemogenetic inhibition of the BLA-vHC circuit decreases anxiety-like behavior on the elevated plus-maze. (A) In rats expressing Gi-DREADD, intra-vHC infusion of CNO (100 ng/side) increased open arm time and open arm entries with no effect on closed arm entries, a measure of locomotor activity (N = 7; *p<0.05, unpaired t test). (B) In rats only expressing the reporter protein, intra-vHC infusion of CNO had no effect on any measures on the elevated plus-maze (N = 5, p >0.05, unpaired t-test).

### 3.2 Chemogenetic inhibition of the BLA-vHC circuit has no effect on ethanol intake

In a separate cohort of rats, we next sought to test the hypothesis that inhibition of the BLA-vHC projection would reduce operant ethanol drinking behaviors. To examine this question, we employed a drinking regimen optimized to separate appetitive and consummatory behaviors, as prior studies by us, and others, have shown that these measures do not correlate with each other and that pharmacological manipulations that inhibit BLA excitability primarily reduce appetitive drinking-related behaviors (Butler *et al*., 2014; McCool *et al*., 2014). In this procedure, rats were trained to complete a 30 lever press response requirement (RR30) to gain access to a sipper tube containing a 10% ethanol solution for daily 20 minute drinking sessions. Ethanol intake/session was used as the primary consummatory measure and averaged 1.27 +/−0.06 g/kg (N=8) in the nine sessions prior to CNO infusions. Tail blood ethanol concentrations were determined on the final drinking session for this cohort and averaged 108.00 +/− 7.94 mg%; N = 8. The number of lever presses during extinction probe trials was used as the main measure of appetitive behavior. In these modified sessions, subjects were allowed to lever press for 20 minute and the ethanol solution was withheld until after this time period had elapsed.

We first examined the effect of bilateral intra-vHC infusions of ACSF and CNO (33, 100, 300 ng/side) in rats expressing hM4D(Gi)-DREADD during standard daily drinking sessions. On microinjection days, sessions were reinforced regardless of whether subjects completed the RR30, as we predicted that intra-vHC CNO infusions might reduce lever pressing. A one-way repeated measures ANOVA, with treatment as the factor, revealed that CNO had no significant effect on ethanol intake (F_3,28_=0.670, p >0.05 (Fig 4A)). All subjects completed the 30 lever press response requirement at the two lowest CNO doses (33, 100 ng/side) but 4 of the 8 subjects did not complete the RR30 at the highest dose tested (300 ng/side). Notably, there were no obvious differences in the effect of CNO on ethanol intake between the subjects that completed the RR30 and those that did not, with CNO reducing intake by 31% in those that completed the response requirement (ACSF= 1.16 +/− 0.05 g/kg; CNO = 0.80 +/− 0.14) and by 21% in those that did not (ACSF= 0.99 +/− 0.15 g/kg; CNO = 0.78 +/− 0.0.09 g/kg). We also tested the highest dose of CNO on a separate cohort of rats that were treated in an identical manner except that they received injections of a viral construct that expressed a reporter protein but not the DREADD. In this group, a one-way repeated measures ANOVA also revealed no effect of treatment on ethanol intake (Fig. 4B). In addition, all subjects completed the 30 lever press response requirement following CNO infusion.

**Figure 4.**
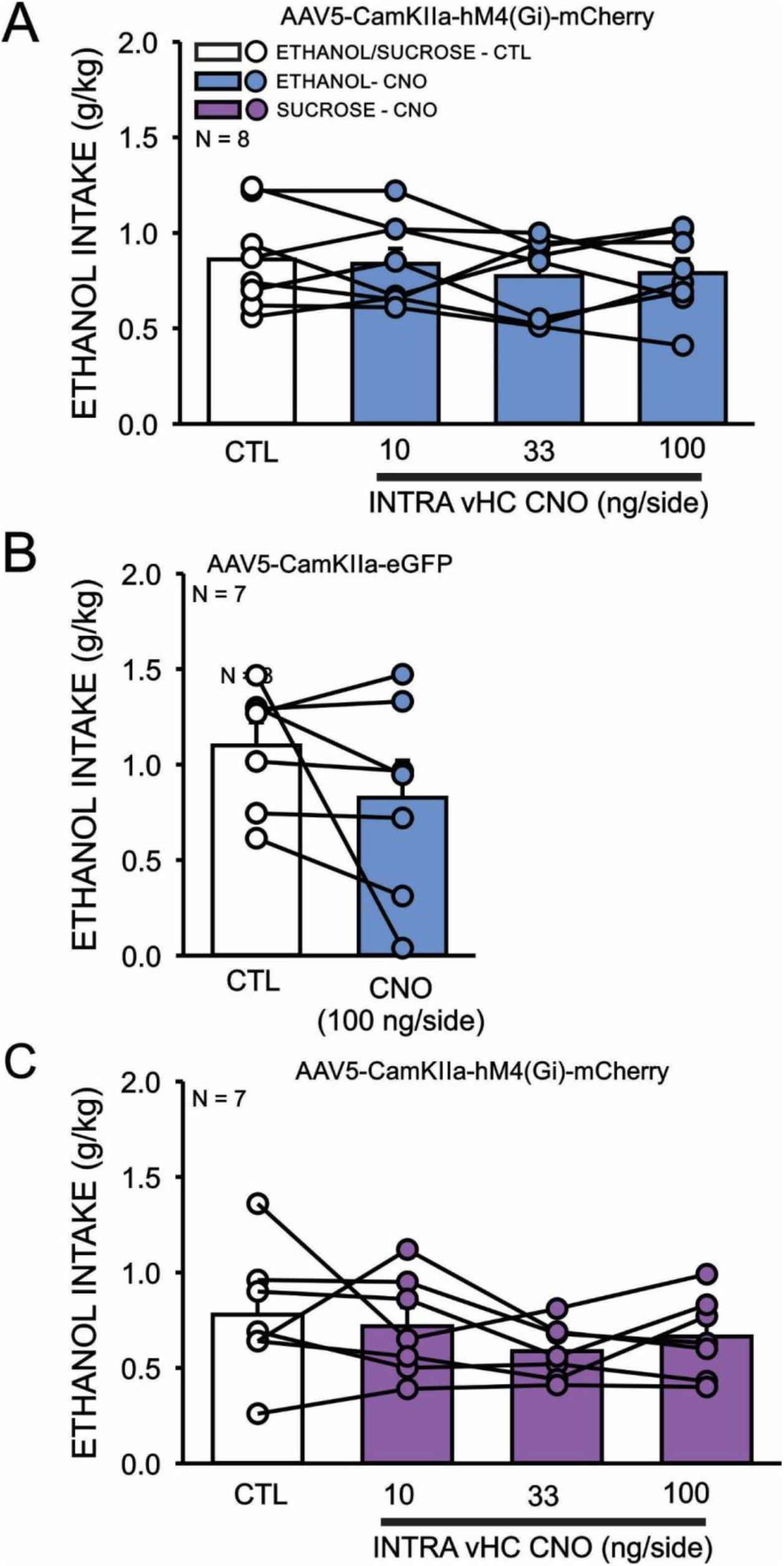
Chemogenetic inhibition of the BLA-vHC circuit has no effect on ethanol or sucrose consumption during operant drinking sessions. (A) In Gi-DREADD expressing rats trained to self-administer 10% ethanol (N = 8), intra-vHC injection of CNO (10, 33, and 100 ng/side) had no effect on ethanol intake during regular operant drinking sessions (one way, repeated measures ANOVA, p > 0.05). (B) Intra-vHC infusion of the reporter protein control group trained to self-administer 10% ethanol (N = 7), intra-vHC infusion of CNO (100 ng/side) had no effect on operant ethanol intake (unpaired t test, p >0.05). (C) In Gi-DREADD expressing rats trained to self-administer 3% sucrose (N = 7), intra-vHC injection of CNO (10, 33, and 100 ng/side) had no effect on operant sucrose intake (one way, repeated measures ANOVA, p > 0.05).

### 3.3 Chemogenetic inhibition of the BLA-vHC circuit has no effect on intake of a natural reinforcer

We also asked whether silencing the BLA-vHC circuit altered intake of a natural reinforcer. To address this question, another cohort of rats was trained to self-administer 3% sucrose using the same limited-access operant procedure used to examine ethanol drinking behaviors. This sucrose concentration engenders similar extinction probe trial responding as 10% ethanol in this operant regimen. A one-way repeated measures ANOVA indicated a significant overall effect of treatment (F_3,24_ = 1.423, p >0.05 (Fig 4C)). All subjects completed the 30 lever press response requirement at the lowest CNO dose (33ng/side) but 2 of the 7 subjects did not complete the RR30 following 100 ng/side CNO and 3 failed to complete after 300 ng/side CNO. As with the ethanol cohorts, CNO did not seem to have a greater effect on sucrose intake in subjects that failed to complete the lever press response requirement. In fact, at the 100 ng/side dose, CNO reduced sucrose intake by 27% in subjects that completed the RR30 (ACSF=0.86 +/− 0.14 g/kg; CNO=0.63 +/− 0.07 g/kg) and by 16% in subjects that did not complete (ACSF=0.58 +/− 0.0.32 g/kg; CNO=0.49 +/−0.08 g/kg). Similarly, 300 ng/side CNO reduced sucrose intake by 17% in rats that completed the RR30 (ACSF=0.90 +/− 0.17 g/kg; CNO=0.74 +/− 0.08 g/kg) and by 11% in subjects that did not complete (ACSF=0.62 +/− 0.19 g/kg; CNO=0.55 +/−0.14 g/kg).

### 3.4 Chemogenetic inhibition of the BLA-vHC circuit reduces extinction probe trial responding for ethanol and sucrose

To test the hypothesis that inhibition of the BLA-vHC pathway would decrease appetitive behaviors, we examined the effect of the intermediate dose of CNO (100 ng/side) on the number of lever presses completed during extinction probe trials, where subjects had 20 minutes to lever press but sessions were not reinforced until the end of this period. The number of lever presses completed in these sessions is a well-validated appetitive measure reflecting a subject’s motivation or craving to obtain the reinforcer (Czachowski *et al*., 2001; Samson & Chappell, 2004; Butler *et al*., 2014). In the ethanol-drinking cohort of rats expressing Gi-DREADD, there was a significant effect of CNO vs ACSF (t_1,14_=2.691, p < 0.02)(Fig 5A).

**Figure 5.**
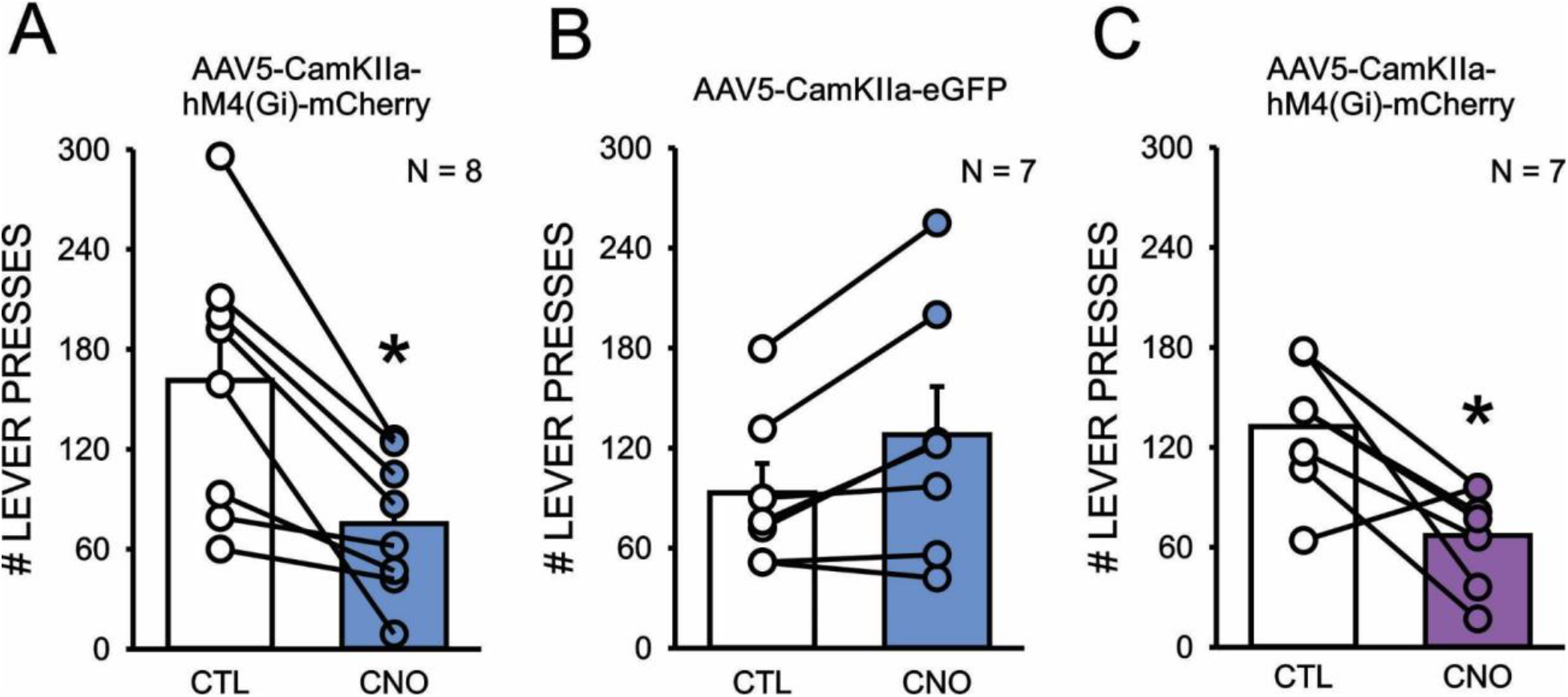
Chemogenetic inhibition of the BLA-vHC reduces extinction probe-trail responding for ethanol and sucrose. (A) Intra-vHC infusion of CNO significantly reduced lever responding during extinction probe trials in Gi-DREADD expressing rats trained to self-administer 10% ethanol (N = 8). (B) Intra-vHC infusion of CNO had no effect on lever responding during extinction probe trials in reporter protein control rats trained to self-administer 10% ethanol (N = 7). (C) Intra-vHC infusion of CNO significantly reduced lever responding during extinction probe trials in Gi-DREADD expressing rats trained to self-administer 3% sucrose (N = 7). *, p <0.05, unpaired t tests.

In contrast, CNO microinfusion had no significant effect on extinction probe trial responding in the cohort that only expressed the reporter protein, confirming that the reduction in this appetitive ethanol drinking behavior was dependent on vHC expression of Gi-DREADD (Fig 5B). Finally, a similar analysis in the sucrose drinking cohort also revealed a significant effect of CNO treatment on extinction probe trial responding (t_1, 12_ = 3.433, p < 0.006). No difference was observed in the magnitude of the effect of intra-vHC CNO on extinction probe trial responding between the ethanol and sucrose cohorts (ethanol = 49.5 +/− 7.7; sucrose = 41.7 +/− 16.7%)(Fig 5C).

### 3.5 Additional comparisons of the effects of chemogenetic silencing of the BLA-vHC circuit on appetitive and consummatory responding for ethanol and sucrose

The preceding analyses revealed that chemogenetic inhibition of the BLA-vHC pathway selectively reduced extinction probe-trial responding, the primary appetitive measure of ethanol drinking-related behavior, for both ethanol and sucrose but this circuit manipulation had no effect on intake of either reinforcer. To gain further insight into the differential effects of this circuit on appetitive and consummatory drinking-related behaviors and further examine possible differences in how this pathway regulates ethanol and sucrose responding, we conducted an additional analysis of three consummatory (latency to first lick, lick rate, number of lick bouts) and three appetitive (latency to first lever press, lever press rate, number of lever press bouts) measures. We examined each of these measures on regular drinking sessions following vehicle or CNO (300 ng/side) and the data were analyzed using a two-way, mixed model ANOVA with reinforcer (ethanol/sucrose) as the between subject factor and treatment (ACSF/CNO) as the within subject factor. In general, this analysis strongly supported our initial findings that inhibiting the BLA-vHC pathway selectively reduced appetitive, but not consummatory, drinking-related behaviors (Fig 6A,B). For example, there was no significant effect of CNO on any of the consummatory measures. In marked contrast, CNO significantly reduced two of the appetitive measures (lever press completion time: F_1,13_ = 24.55, p < 0.001; lever press bouts: F_1,13_ = 10.09, p < 0.01) with a trend of a reduction on latency to first lever press (F1,13 = 3.72, p < 0.08). Notably, there were no significant effects of reinforcer or interactions between this factor and treatment for any of the measures.

**Figure 6.**
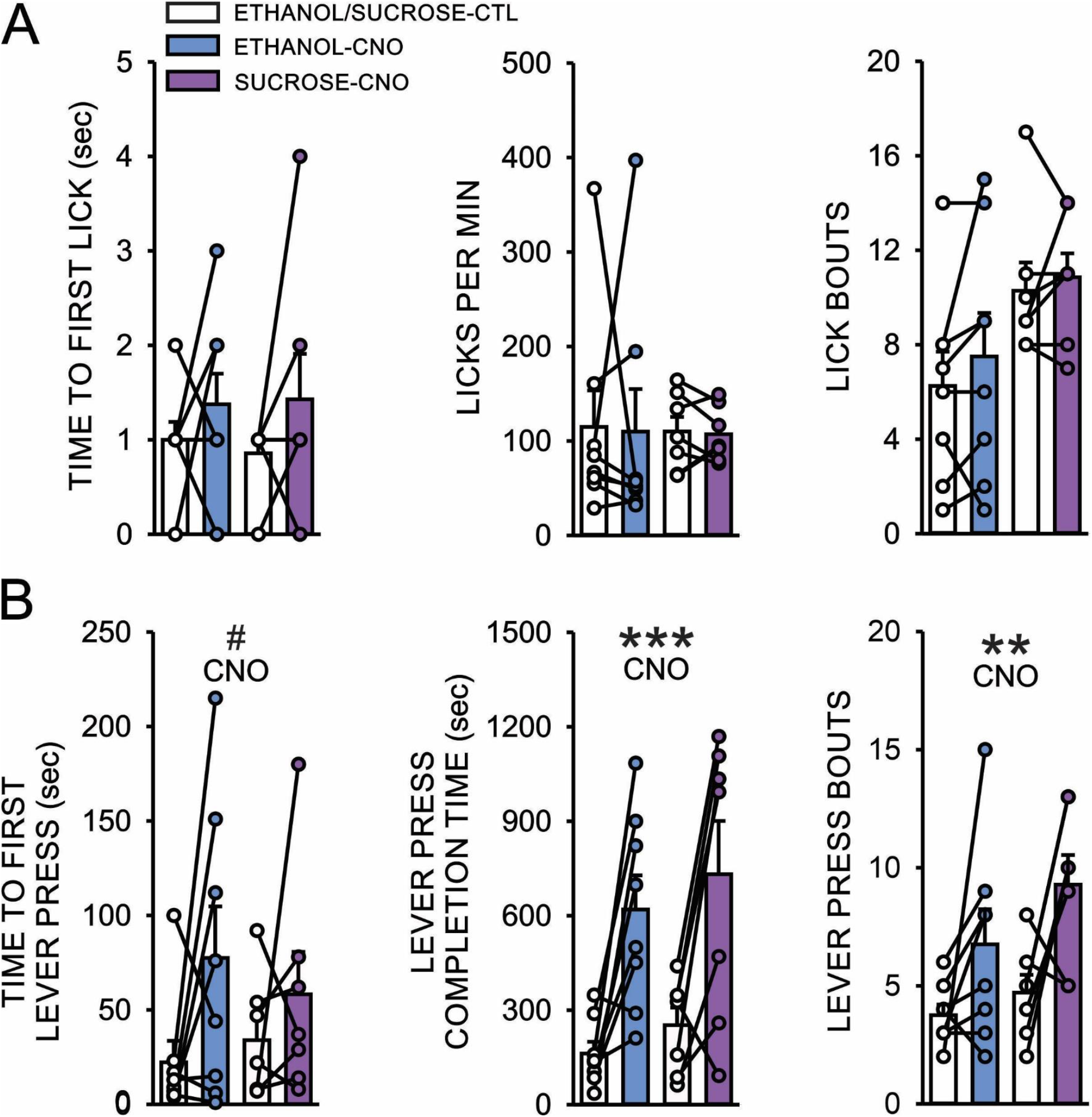
Comparisons of the effects of intra-vHC CNO (300 ng/side) on additional consummatory and appetitive measures, assessed during regular drinking sessions, in ethanol (N = 8) and sucrose (N = 7) groups. A) Chemogenetic inhibition of the BLA-vHC circuit had no effect on any of the consummatory measures (time to first lick, licks/min, lick bouts) in either group. Chemogenetic inhibition of the BLA-vHC circuit significantly reduced two appetitive measures (lever press completion time; lever press bouts with a trend of an inhibition of time to first lever press. No significant effect of reinforcer or interactions between this factor and treatment were observed for any of the consummatory or appetitive measures assessed. **, p < 0.01; ***, p < 0.001, # p < 0.08, mixed model two way ANOVAs.

## 4.0 DISCUSSION

There is compelling evidence, from preclinical and clinical studies, that the etiology of anxiety/stressor disorders and AUD may involve maladaptive changes in common neural circuits. In addition to the frequent co-occurrence of these disorders, withdrawal from chronic alcohol promotes a negative emotional state, of which anxiogenesis is often an integral component (Valdez *et al*., 2002; Morales *et al*., 2015; Morales *et al*., 2018; Ewin *et al*., 2019)

Acute alcohol is a highly effective anxiolytic, even after chronic exposure (McCool *et al*., 2003; Liang *et al*., 2006). Moreover, many medications with anxiolytic properties reduce dependence-induced increases in alcohol drinking in animal models and have shown promise in human AUD clinical trials (e.g. topiramate, baclofen, adrenoceptor modulators) (Silberman *et al*., 2010; de Beaurepaire, 2012; Fox *et al*., 2012; Blodgett *et al*., 2014; Butler *et al*., 2014). Given emerging evidence that a monosynaptic projection from the basolateral amygdala to the ventral hippocampus (BLA-vHC) regulates a broad spectrum of anxiety-related behaviors (Felix-Ortiz *et al*., 2013; Janak & Tye, 2015; Yang *et al*., 2016), here we sought to examine whether this circuit influences operant alcohol drinking measures. We first confirmed that intra-BLA infusion of a viral construct expressing mCherry and Gi-DREADD resulted in robust expression of the reporter protein in the ventral hippocampus and this expression was largely restricted to the CA1 and subiculum. Using ex vivo electrophysiology, we also demonstrated that bath application of CNO significantly inhibited optically-evoked BLA-vHC EPSCs. We then confirmed recent optogenetic findings, demonstrating that intra-vHC infusion of CNO in animals expressing Gi-DREADD within the BLA-vHC circuit significantly decreased anxiety-like behaviors on the elevated plus-maze. Finally, using a limited access, operant drinking regimen, we found that intra-vHC CNO microinfusion selectively inhibited appetitive alcohol drinking-related measures while having no effect on consummatory behaviors. Moreover, this treatment also selectively reduced appetitive behavior for a sucrose reinforcer. Taken together, these findings provide the first evidence in rats that silencing BLA-vHC synaptic communication decreases anxiety-like behaviors as well as appetitive drinking-related behaviors for alcohol and a natural reinforcer (sucrose solution).

We found that chemogenetic inhibition of an excitatory projection from the BLA to the vHC significantly decreased open arm time and open arm entries on the elevated plus-maze with no change in closed arm entries, a measure of nonspecific locomotor activity. Importantly, intra-vHC microinfusion of CNO had no effects on any measures in subjects only expressing a reporter protein in this circuit. These results align with the work of other recent studies that have examined the behavioral role of this circuitry in mice. For example, optogenetic activation of BLA-vHC synapses increased anxiety-like behaviors on the elevated plus-maze while inhibiting this pathway reduced anxiety measures (Felix-Ortiz *et al.*, 2013). Similarly, optogenetic inhibition of this pathway increased center time on the open field and decreased latency to feed on the novelty suppressed feeding task (Felix-Ortiz *et al*., 2013). These studies, along with our current findings, provide further evidence that monosynaptic projections from the BLA to the vHC can bidirectionally modulate a wide range of anxiety-like behaviors.

We then used this same chemogenetic strategy to examine the effect of BLA-vHC circuitry on operant ethanol self-administration using a well-validated regimen that procedurally separates appetitive and consummatory behaviors. During regular drinking sessions, where animals needed to complete a 20 lever press response requirement to gain access to a sipper tube containing 10% ethanol, intra-vHC CNO infusion had no effect on intake. However, during extinction probe trials, where animals had 20 minutes to lever press but the sipper tube was withheld during this period, inhibition of this circuit significantly reduced the number of lever presses completed. This selective effect on appetitive behavior was further supported by examining additional measures on regular drinking sessions. Although intra-vHC CNO had minimal effects on actually completing the lever press response requirement, both lever press completion time and the number of lever press bouts were reduced by inhibiting this circuit whereas consummatory measures, like latency to first lick and lick rate, were unaffected.

These findings are in agreement with prior work by us and others using this same drinking regimen to examine pharmacological manipulations of BLA excitability. Intra-BLA infusion of a serotonin type 2 receptor agonist or a β3-adrenoreceptor agonist, which both elicit inhibition of BLA principal neurons, led to a robust reduction in appetitive measures of ethanol self-administration but had limited effects on consummatory behaviors (Butler *et al*., 2014; McCool *et al*., 2014). Similarly, using a cue-evoked conditioned seeking paradigm, baclofen/muscimol-dependent inhibition of the BLA inhibited ethanol seeking (Chaudhri *et al*.,2013; Millan *et al*., 2015). In contrast to these general manipulations of the BLA, selective activation of a BLA projection to the nucleus accumbens has also been shown to reduce conditioned and unconditioned alcohol consummatory behavior (Millan *et al*., 2017). Taken together these findings indicate that the BLA is involved in both appetitive and consummatory measures of ethanol self-administration but that its dominant influence on these behaviors may be pathway specific. Our data suggests that the BLA-vHC pathway primarily modulates appetitive ethanol drinking-related measures.

Our findings also revealed that BLA-vHC inactivation reduced appetitive measures of sucrose self-administration. These findings suggest that the BLA-vHC circuit may play a more general role in driving motivated behaviors for both natural rewards and drugs of misuse. Indeed, other studies support a role of the vHC in promoting motivated behaviors for natural rewards. For example, increased excitability in a vHC-NAc pathway has been shown to enhance motivation to seek natural rewards (Britt *et al*., 2012; LeGates *et al*., 2018; Reed *et al*.,2018; Gergues *et al*., 2020). Interestingly, activation of the vHC indirectly increases accumbal dopamine release (Zimmerman & Grace, 2016; Lindenbach *et al*., 2019), consistent with its role in promoting motivated behaviors. It should however be noted that, as with the BLA, this brain region can also exert opposite effects on appetitive measures. For example, inhibition of the ventral hippocampus has been found to actually play a permissive role in goal-directed behavior to obtain natural rewards (Kosugi *et al*., 2021; Yoshida *et al*., 2021)

Although inhibition of the BLA-vHC circuit had similar effects on ethanol and sucrose seeking-related measures, it is important to note that these studies were conducted in rats that were not physically dependent on ethanol. It will be important, in future studies, to conduct a similar investigation in ethanol-dependent animals to determine if the negative emotional state that develops during withdrawal differentially influences how this pathway regulates seeking for ethanol and natural reinforcers. The vHC has also been recognized for its role as a gatekeeper for changes in behavioral contingencies. Activation of the vHC is thought to block the acquisition of novel approaches to shifting behavioral paradigms and thus maintain habituated responses (Barker *et al*., 2019). Our operant paradigm may produce habituated responses that are protected by the vHC’s role in inhibiting changes in behavioral contingencies, as experienced by our animals during extinction (where the completion of the habituated number of lever presses needed to receive the reward does not lead to its delivery). Thus, the reduction in the number of completed lever presses through CNO-dependent inhibition of the BLA-vHC circuit could be explained by a disinhibition of contingency updating. However, we also observed a reduction in the “response requirement” completion time during regular drinking sessions in response to chemogenetic BLA-vHC circuit inhibition. In this condition, animals should not yet have detected any change in contingency and therefore a failure in contingency updating is unlikely to be the sole driver in the reduction of appetitive measures we observed.

Another important consideration is the dichotomy of valence information that is processed by the BLA. Although there is now general agreement that sensory and emotional stimuli with positive and negative valance are encoded by distinct populations of BLA pyramidal neurons, how these cells are anatomically segregated remains an active area of debate. While some studies have shown that positive and negative valence-encoding neurons are intermingled throughout the BLA, others have argued for regional segregation along the anterior-posterior axis (Kim *et al*., 2016; Beyeler *et al*., 2018; Wang *et al*., 2018; Pi *et al*., 2020). Based on our histological analysis, we identified no biased anterior or posterior regional expression, with viral expression indicating a relatively even distribution throughout the BLA. Furthermore, the viral vector promoter we used (CamKII) is thought to predominantly target glutamatergic projection neurons in a non-specific manner. Taken together, our experimental approaches argue against us having biased inhibition in the BLA-vHC pathway towards any known positively or negatively valanced information processing.

One limitation of our study is that we exclusively used male rats, although similar studies using females are underway. Prior findings from our lab have identified profound sex differences in vHC synaptic adaptations promoted by chronic intermittent ethanol exposure (CIE), a rodent model of alcohol dependence. Although withdrawal from CIE was associated with increased anxiety-like behavior in both sexes, this adaptation was accompanied by vHC hyperexcitability in male rats and decreased vHC network excitability in females (Ewin *et al*., 2019; Bach *et al*.,2021a; Bach *et al*., 2021b). Additional studies will be needed to determine how these sexually dimorphic adaptations associated with chronic ethanol exposure relate to motivational (appetitive) aspects of drinking in male and female rats. Although it would seem unlikely that there are sex differences in the circuitry that governs appetitive behaviors, a recent set of studies reported that chemogenetic inhibition of the NAc had opposite effects on ethanol drinking in male and female mice (Purohit *et al*., 2018; Townsley *et al*., 2021).

In summary, our findings confirm prior work in mice, that an excitatory projection from the BLA to the vHC modulates anxiety-like behavior in outbred rats. Using an operant paradigm that can distinguish between appetitive and consummatory behaviors we also demonstrated, for the first time, that inhibition of this pathway selectively reduces seeking-related behaviors for ethanol and a natural reward. These findings, along with our recent discovery that withdrawal from chronic ethanol increases excitability in this circuit (Bach *et al*., 2021a) are consistent with the idea that maladaptive increases in BLA-vHC synaptic communication may contribute to some of the behavioral symptoms associated with AUD and comorbid anxiety disorders. Our data also suggest that pharmacological interventions that reduce BLA-vHC excitation may be particularly effective treatments for individuals suffering from these disorders.

## Conflict of Interest

The authors declare no competing financial interests.

## Acknowledgements

This work was supported by the NIH/NIAAA awards: AA25819 (SEE), AA7565 (AGA), AA29292 (HNC), AA26117, AA17531, AA26455 and AA26551 (JLW).

## Notes

### Competing Interest Statement

The authors have declared no competing interest.

### Summary of Updates

This manuscript is currently under review at a new journal and has been substantively revised. In addition, a new author, Hannah N. Carlson, who made important contributions to the revised manuscript, has been added as an author.

